# An investigation into serotonergic and environmental interventions against depression in a simulated delayed reward paradigm

**DOI:** 10.1101/580456

**Authors:** Bernd Porr, Alex Trew, Alice Miller

## Abstract

The disruption of the serotonergic (5HT) system has been implicated in causing major depression and the standard view is that a lack of serotonin is to blame for the resulting symptoms. Consequently, pharmacological interventions aim to increase serotonin concentration in its target areas or stimulating excitatory 5HT receptors. A standard approach is to use serotonin reuptake inhibitors (SSRIs) which cause a higher accumulation of serotonin. Another approach is to stimulate excitatory serotonin receptors with psychedelic drugs. This paper compares these two approaches by first setting up a system level limbic system model of the relevant brain areas and then modelling a delayed reward paradigm which is known to be disrupted by a lack of 5HT. Central to our model is how serotonin changes the response characteristics of decision making neurons where low levels of 5HT allows small signals to pass through whereas high levels of 5HT create a barrier for smaller signals but amplifying larger ones. We show with both standard behavioural simulations and model checking that SSRIs perform significantly better against interventions with psychedelics. However, psychedelics might work better in other paradigms where a high level of exploration is beneficial to obtain rewards.

## 1 Introduction

Serotonin (5HT) has been implicated in causing major mood disorders such as depression (Chaudhury et al., 2015). Consequently, influencing the serotonergic system with pharmacological interventions has been shown to be effective. In particular, serotonin reuptake inhibitors (SSRIs) have positive effects on a patient’s mood (Barker and Blakely, 1995; Cipriani et al., 2012; Stahl, 1994). However, this well established therapy has its critics who favour psychedelics instead of SSRIs as a drug for combating depression (Carhart-Harris et al., 2014). Psychedelics are mainly known for their ability to alter the perception of sensor stimuli as shown with drugs such as LSD (Winter, 2009). In addition SSRIs change the perception of emotionally related stimuli so could be used to indirectly influence a subject’s mood (Harmer, 2008). The fact that 5HT can alter cortical processing suggests that the role of the serotonin receptors should be investigated more closely (Mengod et al., 2009). While there are over 7 different 5HT receptors, 5HTR1 and 5HTR2 have been mainly implicated in mood disorders (Carhart-Harris and Nutt, 2017). This means that to understand the action of 5HT we must at least determine how 5HTR1 and 5HTR2 operate together to influence neuronal processing.

However, mood is not just linked to serotonin but also to dopamine (Schultz, 1998; Cofer, 1981). This means serotonin cannot be understood in isolation but must be considered in conjunction with the dopaminergic system (Schildkraut, 1965; Martin-Soelch, 2009). In the past dopamine was prominent in models of reward based processing and 5HT was just seen as an inverted version of the dopaminergic signal (Boureau and Dayan, 2011). However this view has been abandoned (Dayan and Huys, 2015) in the light of recent experimental results which show that 5HT tracks the long term anticipation of a reward (Nakamura et al., 2008; Li et al., 2016). Thus, serotonin codes distinctly different information to dopamine.

What kind of behaviour is improved by the release of serotonin? From recent studies it has become apparent that serotonin is required for situations where an animal needs to wait to obtain a delayed reward (Li et al., 2016; Bari and Robbins, 2013) and that 5HT “integrates expected, or changes in, relevant sensory and emotional internal/external information” (Homberg, 2012).

To understand the role of 5HT we need to investigate the following:

- the action of 5HT on the major 5HT receptors, in particular 5HTR1 and 5HTR2
- processing of emotionally relevant stimuli from sensor to action via both cortical and subcortical structures
- how processing/perception of these stimuli is altered/controlled by 5HT
- earmarking behavioural paradigms which require a functional 5HT system
- how the reward system is impacted by altered 5HT signal processing

In other words: we claim that it is only possible to understand the 5HT system by using a holistic approach including all levels starting from 5HT receptors up to behaviour. This means that possible interventions cannot be seen in isolation but need to be viewed in combination.

The standard approach of testing a system model such as ours is to run simulations of the model many times for each proposed set of parameters and perform statistical analysis on the results in each case. This has the advantage that both behaviour and system can be modelled at great detail. However, running multiple simulations is time-consuming and does not guarantee complete coverage (how many simulations should we run?) In this paper we use an additional approach: Model Checking. This is a technique in which a system is expressed using a formal language and converted into finite state model. This underlying model can then be used to exhaustively check properties (e.g. the probability of an event occurring in the long run) for a range of parameter values. In particular here we will use model checking to investigate the link between the neuronal response characteristic and its impact on delayed reward learning. Both the behavioural simulator and the model are available via an open access repository (Porr et al., 2019).

The paper is structured as follows: first, we present a behavioural experiment which involves a rat waiting for a delayed reward. Then we describe the information flow from sensor inputs to actions via cortical and sub-cortical structures. We then focus on how information processing is altered with the action of serotonin (5HT), how sensor inputs are processed differently depending on the concentration of 5HT and how this is achieved with the two 5HT receptors 5HTR1/5HTR2. Finally the model is completed by adding the dopaminergic reward system. We then run simulations where 5HT is reduced and different interventions such as SSRIs, psychedelics and environmental changes are introduced so that the rat receives more rewards. We will then draw our conclusions as to which of these interventions are successful and under which conditions.

## 2 Methods

### 2.1 The behavioural experiment: patience to obtain a reward

Fig. 1 illustrates our experiment, which can loosely be described as “having patience to receive a reward” (Li et al., 2016). A rat needs to learn to approach the green landmark on the left and then wait there until food becomes available and thus visible. The blue landmark on the right is for distraction: it will never show any reward but it generates visual information. We can divide the learning behaviour into five steps:

1. the rat accidentally waits in front of the landmark and receives the reward
2. next time the rat has associated the visual information of the landmark with the reward and approaches it.
3. at the same it has also associated the area around the landmark as a place to wait
4. the food appears and the rat approaches the food
5. (as in 1) the food results in a reward

**Figure 1:**
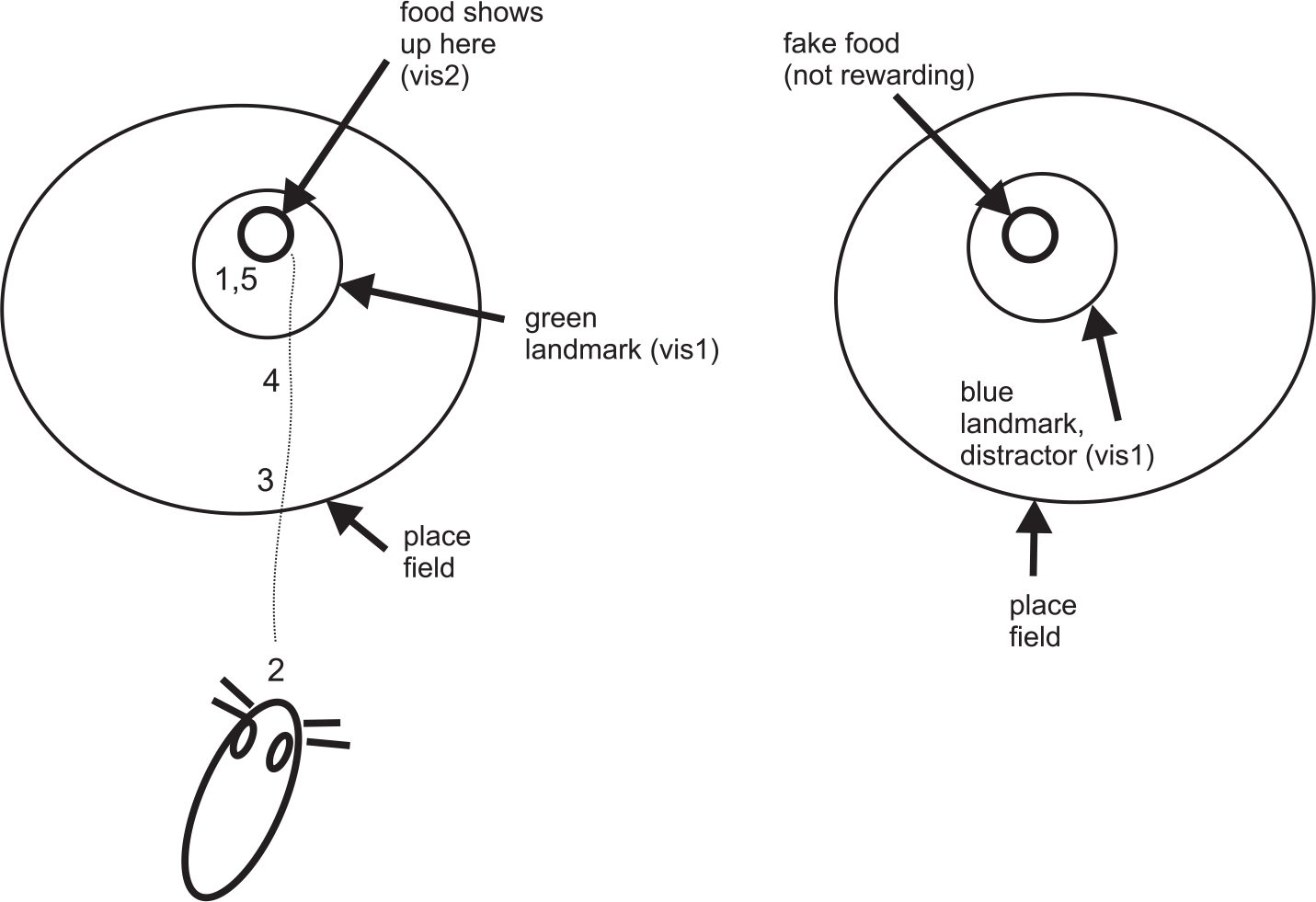
Behavioural experiment

These steps emerge naturally just by walking through the behaviour and our task now is to identify neuronal structures which generate this behaviour.

### 2.2 From sensor to action: the limbic system model

By building our limbic system model we first need to focus on those nuclei which translate a pre-processed sensor signal into an action, thus enabling our simulated rat to approach a landmark which eventually releases food. We will then expand it to the complete model.

#### 2.2.1 Action selection in the mPFC and NAcc

We first describe how a sensor signal causes an action and how this processing is modulated by both serotonin and dopamine. Fig. 2 shows the relevant nuclei: medial prefrontal cortex (mPFC) and Nucleus Accumbens (NAcc) (Berthoud, 2004). The mPFC receives excitatory inputs from primary sensor areas such as visual, smell and tactile. In addition it might also receive input from the hippocampus and other higher level areas which are strongly linked to sensor information and context. These inputs are generally glutamatergic (GLU) and excite the neurons in the mPFC. The prominent neuromodulator here is serotonin (5HT) which is released into the mPFC and other cortical areas from the dorsal Raphe nucleus (DRN) (Linley et al., 2013). A major output target of the mPFC is the NAcc, in particular the NAcc core (Heimer et al., 1991; Brog et al., 1993). The NAcc core is closely related to the more dorsal areas of the striatum and is responsible for action selection. Synapses here are strongly modulated by dopamine. The output of the NAcc core then triggers motor actions via a polysynaptic pathway which targets the motor cortices (Kelley, 2004; Humphries and Prescott, 2010). In our example these are just two actions: approach the green landmark or approach the blue one. Of course a real animal has more pathways but we focus on two processing streams which are sufficient for our simple experiment. Because the sensor signals progress from the sensor areas through the mPFC and then the NAcc core, we first describe the mPFC and then the NAcc core.

**Figure 2:**
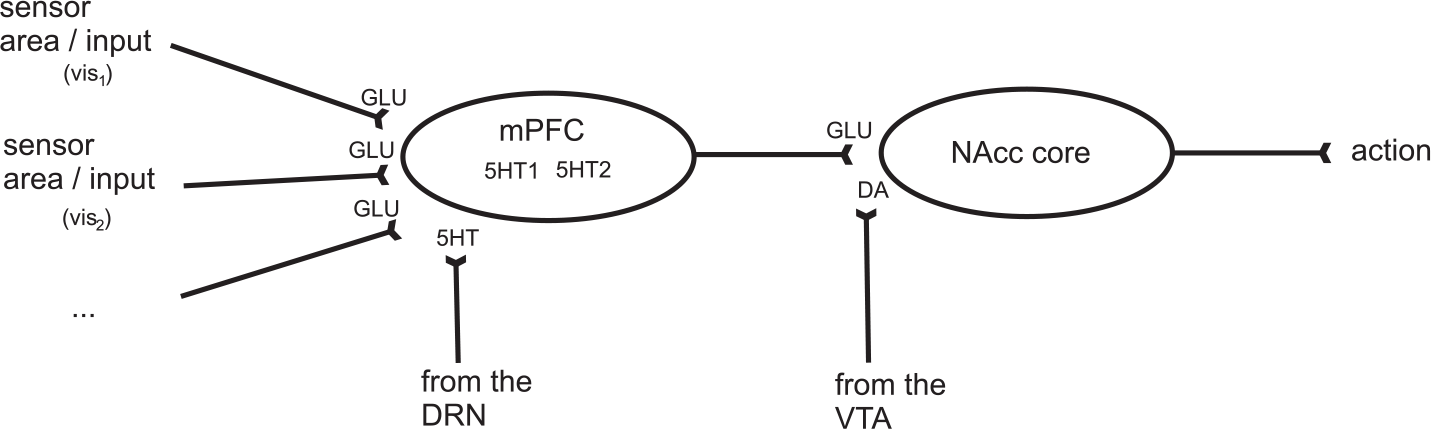
The signal flow from the cortex (mPFC) to the nucleus accumbens (NAcc core)

#### 2.2.2 The action of 5HTR1 and 5HTR2 receptors in the mPFC

As outlined above the mPFC integrates information from numerous primary and secondary sensor areas but the important aspect is that it receives a strong serotonergic innervation. Serotonergic fibers originate in the dorsal Raphe Nucleus (DRN) and from there they mainly target prefrontal cortical areas and to a lesser extent primary sensor areas and subcortical areas (Linley et al., 2013). However, we simply focus on the strongest innervations of 5HT and these occur in the prefrontal areas. There are two major receptors in the cortex: 5HTR1 and 5HTR2 (Palacios et al., 1990; Mengod et al., 2009). While 5HTR1 is inhibitory, 5HTR2 is excitatory. Recent findings point to a more sophisticated role of these receptors when investigating response curves of cortical processing as done in the visual cortex (Shimegi et al., 2016; Seillier et al., 2017) and we propose that 5HT alters the signal processing in the mPFC in a similar way.

##### 5HTR1

Based on Shimegi et al. (2016) we model the action of the receptor 5HTR1 as a parameter in a psychometric function which slowly saturates towards one:

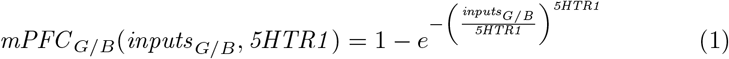

where *inputs*_*G/B*_ is the sum of inputs to the mPFC(see Fig. 2)

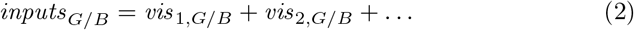

*mPFC_G/B_* is the output of the mPFC and *5HTR1* the activation of the 5HTR1 receptor. The subscripts “G/B” indicate that these are two pathways through the mPFC: one to target the green landmark and one to target the blue one. Fig. 3A shows the response of an mPFC neuron at different 5HTR1 activations (1,2,3). At low 5HTR1 activations (*5HTR1* = 1) low cortical inputs (*inputs* < 1.5) are amplified whereas when the 5HTR1 activation is high (*5HTR1* > 2) lower cortical input values (*inputs* < 1.5) are suppressed. This means that when 5HT is small signals are amplified which in turn makes the animal very attentive to small/noisy cues. On the other hand if 5HT is high lower input signals to the cortex are suppressed. Weak cues or any kind of distraction will be suppressed whereas strong stimuli will be more amplified.

**Figure 3:**
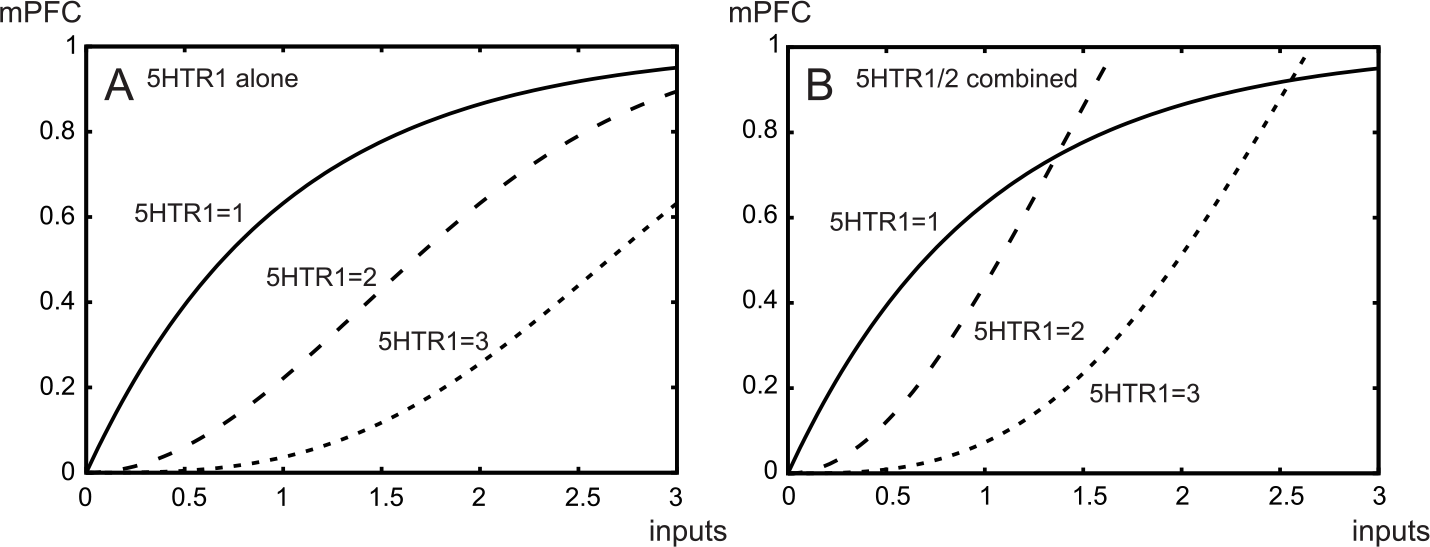
The response functions of mPFC neurons: A) altered by 5HTR1 alone and B) by 5HTR1/R2 combined.

##### 5HTR2

The action of the receptor 5HTR2 can be formulated in a much simpler way: it adds a certain *gain* to the processing in a cortical neuron which can be seen as a multiplicative term which then scales Eq. 1 and reflects the model by Carhart-Harris and Nutt (2017) and the findings by Shimegi et al. (2016). It can be seen in Fig. 3B that the effect of the gain is seen mainly for strong cortical inputs and these are then disproportionally amplified. This means that strong cortical inputs receive an additional boost and might be executed with a strong *vigour* Cofer and Appley (1964).

##### Combined action of 5HTR1 and 5HTR2 in the mPFC

We can now combine the action of both serotonin receptors which results in the following equation describing how serotonin influences cortical processing:

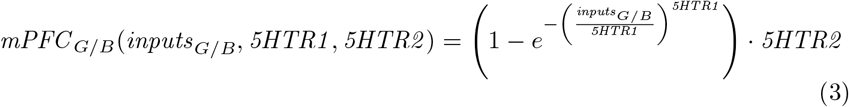

where *inputs_G/B_* and *mPFC_G/B_* are the total input and output to/from the mPFC respectively. In our example we have two pathways to consider: one for the green landmark and one for the blue.

We need to establish a relationship between the 5HT concentration and the activation of the receptors *5HTR1* and *5HTR2*. In general this means that we have a mapping from the 5HT concentration to the receptor activation which we assume to be linear:

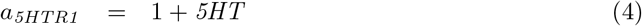

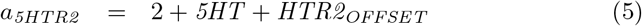

where adding the constants 1 and 2 for *a*_*5HTR1*_ and *a*_*5HTR2*_ respectively guarantees a baseline throughput of the signals in Eq. 3 so that a signal passes through even when *DRN* = 0.

The constant *HTR2*_*OFFSET*_ is usually zero but is set to a non-zero value when we want to simulate the effect of psychedelics. This will allow us to investigate whether psychedelics are able to reverse the deficit caused by excessive 5HT inhibition.

Overall this means that with high 5HT concentrations inputs to cortical circuits need to have a high salience to coincide with other inputs. For example the visual cue of a food dispenser needs to coincide with the visual input of the food itself at the moment it is delivered. This means that the cortex is both an integrator of information and a gatekeeper. It transmits information to the decision making circuitry in the NAcc core.

#### 2.2.3 Reward based learning in the NAcc core

So far we have shaped the signals in terms of attention or signal to noise but have not associated it with any reward. The mPFC projects to the NAcc core which receives a strong dopaminergic (DA) modulation. This has a certain baseline concentration and can either increase or decrease causing a corresponding increase or decrease in synaptic strength of the mPFC input (Beckstead et al., 1979; Humphries and Prescott, 2010):

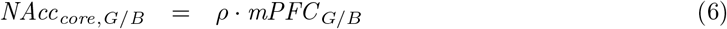

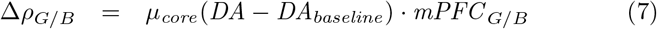

where *ρ*_*G/B*_ are the weights of the projections from the mPFC to the NAcc core, *DA* the dopamine in in the NAcc core released from the ventral tegmental area (VTA) and *DA*_*baseline*_ the baseline DA concentration. This means that a DA concentration above or below baseline corresponds to an occurrence of long term potentiation (LTP) or long term depression (LTD) respectively. This implements weight changes which are compatible with the classical reward prediction error (Schultz et al., 1997). If a reward is encountered unexpectedly the DA concentration increases and if a reward is omitted unexpectedly the DA concentration decreases (referred to as the “dip”).

Note that the cortex also receives a DA modulation (Beckstead et al., 1979) and the NAcc 5HT modulation (Vertes et al., 2010). However, these effects are small in contrast to cortical 5HT modulation and NAcc DA modulation. Thus, to keep the model clear and distinct we broadly state that the cortex is modulated via 5HT while the subcortical areas perform reinforcement learning via DA.

As a final step we add the circuitry which computes both the 5HT and DA activity which are released from the ventral tegmental area (VTA) (Beckstead et al., 1979) and the dorsal Raphe nucleus (DRN) respectively. This leads to the complete limbic system model.

### 2.3 Complete circuit model

So far we have dealt only with the novel aspects of our limbic system model which explains how a variable response curve in the cortex and learning in the NAcc core leads to behaviour associated with waiting for a delayed reward. However, we also require the circuitry which generates the signals for both the VTA and the DRN. While the DRN becomes active in anticipation of a reward, the VTA exhibits a classical error signal which only becomes active when the reward is unexpected, and then its amplitude slowly decays. We need to describe how these two signals are computed and the corresponding circuit is shown in Fig. 4.

**Figure 4:**
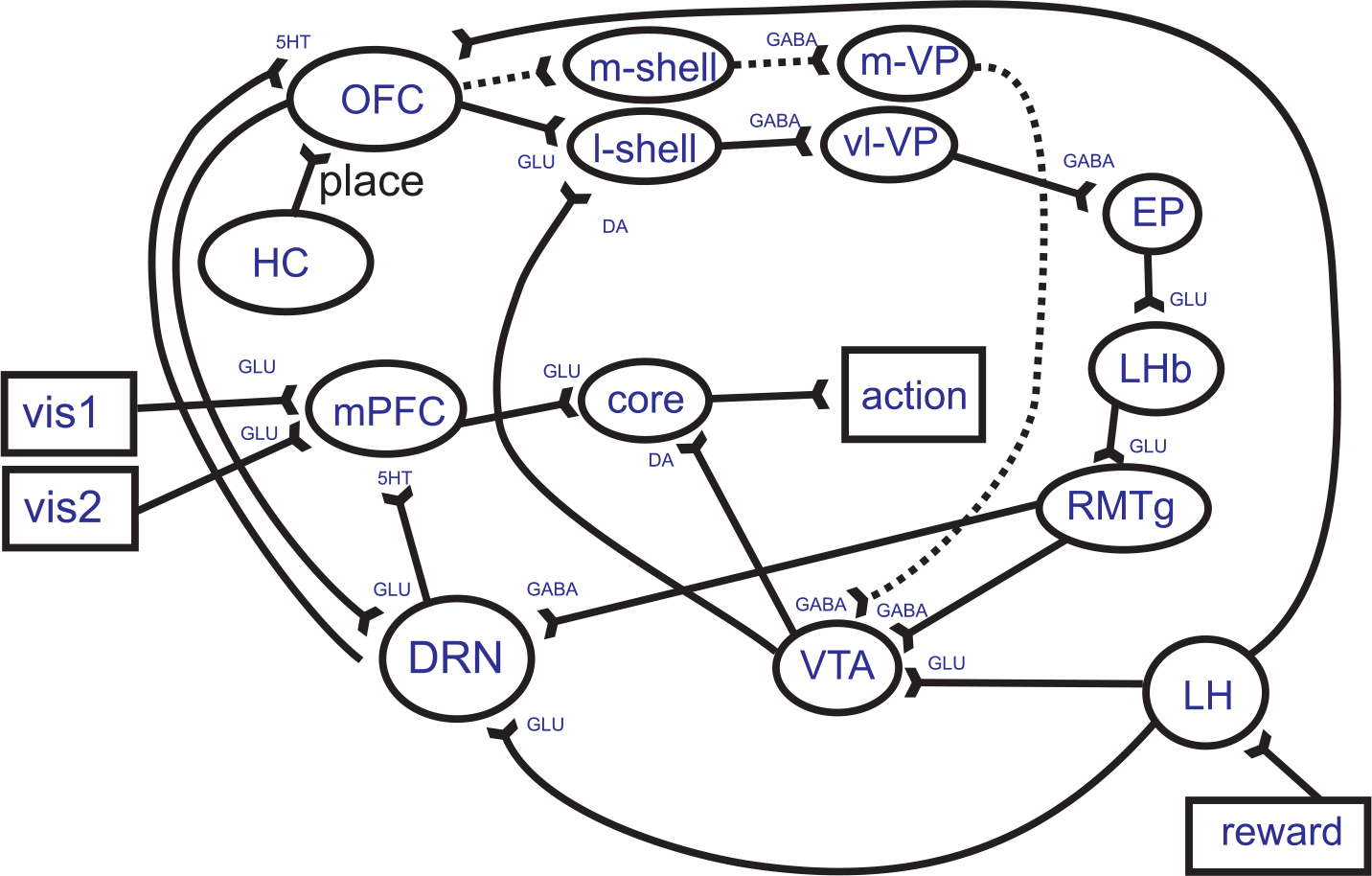
Full limbic circuit

#### 2.3.1 VTA

We start with the activity in the ventral tegmental area (VTA) (Sesack and Grace, 2010). A direct pathway from the lateral hypothalamus (LH) to the VTA drives the VTA whenever a primary reward has been encountered. The lateral hypothalamus is well known to respond to primary rewards. However it is also well known that once the reward can be predicted the activity in the VTA will diminish. This is achieved by the pathway: OFC – l-shell – vl-VP – EP – LHb – RMTg and then VTA. Overall this path is inhibitory. The OFC and the NAcc l-shell learn to associate cues with the primary reward which in turn inhibit the VTA. In addition cues or conditioned stimuli cause bursts in the VTA which are conveyed via the m-shell – m-VP – VTA pathway. However, this pathway is not modelled as we do not need second order conditioning here.

#### 2.3.2 DRN

The main focus of this paper is about serotonin (5HT) which is mainly released from neurons in the dorsal Raphe nucleus (DRN) (Michelsen et al., 2007; Pollak Dorocic et al., 2014). The DRN receives an excitatory input from the lateral hypothalamus (LH) (Aghajanian et al., 1990; Lee et al., 2003) which becomes active when a primary reward is experienced. Again, as with the VTA the signal in the DRN diminishes via the slowly increasing activity in the RMTg–DRN pathway. However, the main difference to the VTA is *the intimate reciprocal connection to the prefrontal cortex* (Zhou et al., 2015; Roberts, 2011), in particular we are interested in the orbitofrontal cortex (OFC). Apart from the input from the LH this is the main excitatory input to the DRN (Zhou et al., 2015). We propose that the sustained activity of the DRN in anticipation of a reward is solely generated by cortical structures and in particular by the OFC.

As mentioned above the OFC learns to associate stimuli with the reward. These could be direct sensor inputs or pre-processed ones. In our case we assume that the OFC receives place information from the hippocampus and can then “remember” that a reward has occurred at that place. Of course the OFC has many additional abilities. In particular for reversal learning to provide persistent activity which lasts after a reward has been omitted and can provide long lasting depression of both VTA and DRN neurons via the RMTg. However, this is beyond the scope of this paper in which we just focus on reward acquisition.

This leads us to the following equations to calculate the activity of the DRN. Since the OFC projects into the DRN we need to define its activity first:

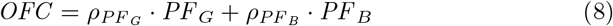

where *PF*_*G*_, *PF*_*B*_ are hippocampal place fields around the green and blue land-marks respectively and 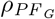, 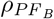 the weights feeding these place fields into the OFC. The two weights from the hippocampus to the OFC change according to:

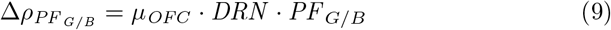

where the activity of the DRN is calculated as:

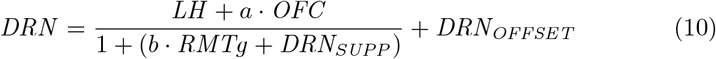

and *LH* is the activity of the lateral hypothalamus (LH) which becomes active when encountering a primary reward. The *RMTg* provides a negative feedback on the OFC via the same subcortical pathway as for the reward prediction error and *a*, *b* are scaling constants.

To test the pathological cases we have introduced two constants: *DRN*_*SUPP*_ is zero under control conditions and set to non-zero to simulate excessive tonic inhibition for pathological DRN hypoactivity. Similarly *DRN*_*OFFSET*_ is zero for control but will be set to a positive value to simulate the effect of the serotonin re-uptake inhibitor.

When does the DRN become active? Consider Eq. 10 that shows how the DRN fires via the LH pathway at the moment a reward appears. We propose that 5HT causes learning in the OFC and associates the place field with the reward. Note that it is likely that a small VTA innervation will cause plasticity in the OFC to be increased. However we separate the roles of 5HT and DA between cortical and subcortical processing and propose that plasticity in the OFC is triggered by 5HT (Peñas-Cazorla and Vilaró, 2015; Roberts, 2011; Mlinar et al., 2006; Phillips et al., 2018).

Before we run simulations we examine graphical traces of the relevant signals to prepare for the more complex signals in the real simulation run.

### 2.4 Linking behaviour to the signals

How is the behaviour of the rat in our experiment linked to the neuronal model described above? Before stating the equations we go through the activity with the help of the traces in Fig. 5 which represent a cortical mPFC neuron processing approach behaviour to the left reward site which will provide delayed rewards.

1. When the rat encounters the primary reward at the green landmark the LH fires which in turn makes the NAcc core learn to associate the visual information of the landmark *vis*_1,*G*_ with the reward. This will guarantee that the rat will approach the green landmark from this distance. At the same time the primary reward is transmitted from the LH to the OFC which associates the place field (circle around the landmark) with the reward.
2. The rat sees the landmark from a distance so input *vis*_1,*G*_ is active. The food is not yet shown so *vis*_2,*G*_ is zero. The activity of 5HT is zero which allows the activity *vis*_1,*G*_ to progress easily via the mPFC and NAcc core to the motor circuits, thus causing the rat to approach the landmark.
3. The rat enters the place field. The hippocampus now provides place field information to the OFC which in turn drives the DRN. This means that the 5HT is released in the mPFC where it changes the signal to noise ratio of the incoming signals. Recall that at this point *vis*_1,*G*_ is greater than zero (indicating “go to landmark”) whereas *vis*_2,*G*_ is zero. This will effectively make the output of the mPFC smaller which can be seen in the cartoon.
4. After a delay the food appears and *vis*_2,*G*_ > 0 which, in conjunction with *vis*_1,*G*_ = 0, results in a strong input to the mPFC which can progress to the NAcc core and cause the animal to approach the food.

**Figure 5:**
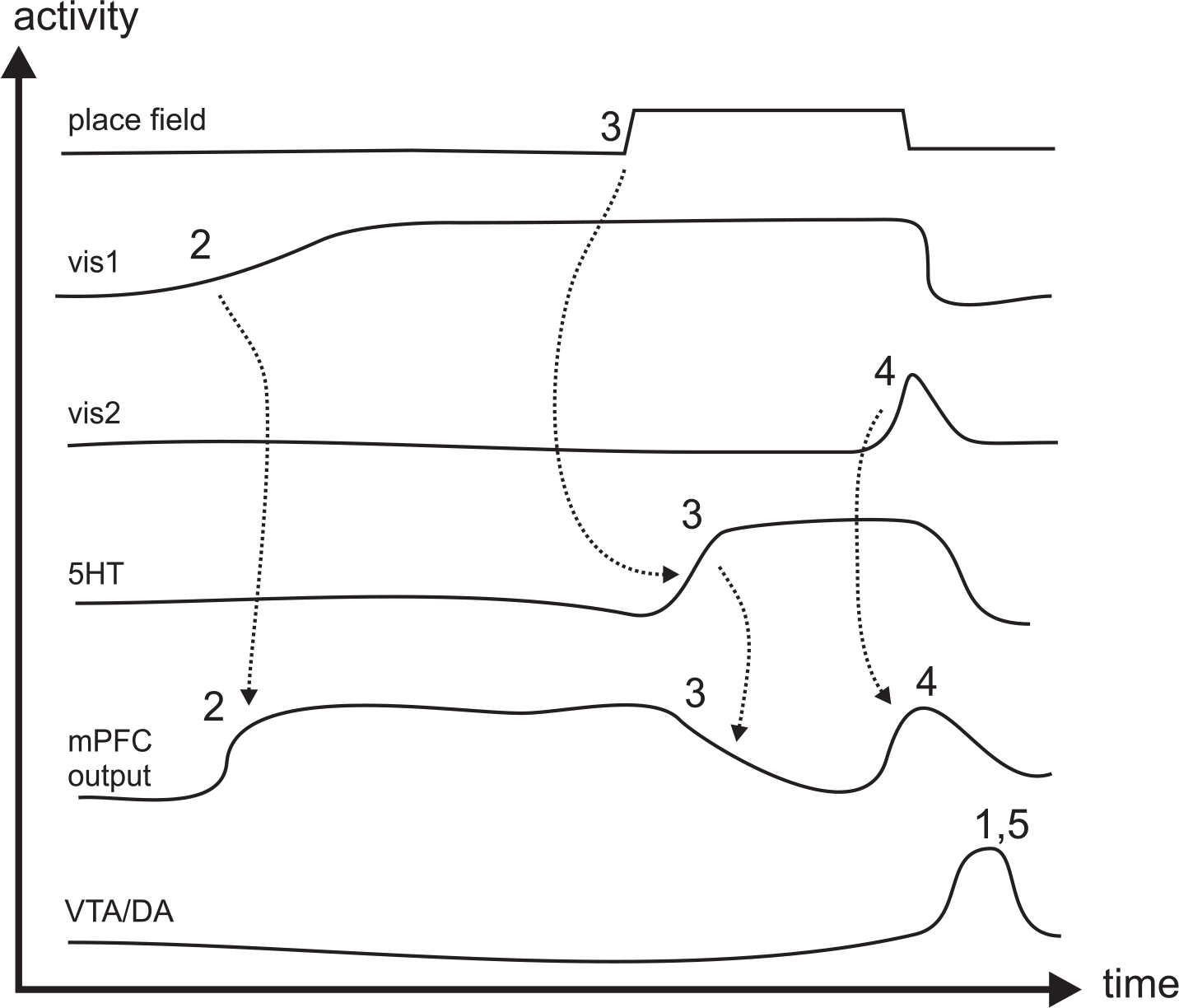
Activity cartoon traces

In summary we have described a model of a decision-making network that spans cortical and subcortical areas. The cortex shapes the signals so that with low 5HT concentrations small stimuli cause an action whereas with high 5HT concentrations only strong or combined stimuli can progress to the NAcc core to trigger actions. The subcortical areas are therefore responsible for reinforcement learning.

### 2.5 Scenarios to investigate drug action

Central to this paper are models against depression. It is widely accepted that a hypofunction of the DRN causes less 5HT to be released. There are several proposed solutions to this problem and we will investigate them in this paper. Our aim is to determine which of the proposed solutions will indeed prove to be beneficial, and which will be shown to be counterproductive. To do this, after a successful run of a healthy animal we will reduce the release of 5HT by disrupting DRN activation both directly and indirectly by increasing the activity of the lateral Habenula. We will observe the behavioural effects and measure the impact of different interventions.

The scenarios for the statistical analysis have an additional parameter which complements the pathological interventions above: the time the agent must wait for a reward. The default number of time steps is 150. By reducing this period to 100 steps in some cases we can model two interventions:

1. Pharmacological intervention: SSRIs or 5HT receptor agonists such as LSD or magic mushrooms.
2. Environmental intervention: the time the agent needs to wait till it receives its reward.

As we have a scenario where waiting is crucial to obtaining a reward a reduced waiting time is the obvious intervention here. This can then be compared to the pharmacological intervention in terms of its effect.

We combine our (two pharmacological and one environmental) interventions to allow us to consider the following scenarios. In all cases the reward is delayed the default number of time steps unless a reduced reward delay is indicated:

1. Control: all parameters were based on those from Section 3.1.1 where the simulated rat successfully waits in front of the landmark.
2. Reward early: same parameters as above and as in Section 3.1.1 but the food appears earlier (reduced reward delay).
3. DRN suppress: the DRN activity is suppressed by a GABAergic efference (*DRN_SUPP_* > 0 in Eq. 10).
4. DRN suppress & reward early: the parameters are the same as in 3 but with a reduced reward delay.
5. DRN suppress & SSRI: this matches Section 3.1.3 where the SSRIs cause a constant baseline shift of the 5HT receptor activations because of slow 5HT reuptake (*DRN_SUPP_* > 0, *DRN*_*OFFSET*_ > 0 in Eq. 10).
6. DRN suppress & SSRI & reward early: this analysis has the same parameters as 5) but with a reduced reward delay.
7. DRN suppress & 5HTR2 agonist: the DRN activity is suppressed by a GABAergic efference (*DRN*_*SUPP*_ > 0 in Eq. 10) and the 5HTR2 receptor is tonically stimulated (*HTR2*_*OFFSET*_ > 0, Eq. 5) so that the gain of the transmission is increased as described in Section 3.1.4.
8. Same parameters as in 7 but with a reduced reward delay.

These different scenarios can be investigated both in single runs to gain a deep understanding of the interactions between the nuclei, and by conducting multiple random runs to determine statistics indicating how successful learning has been. For the behaviour based approach we conduct *Monte Carlo* based experiments for each scenario to calculate the relative reward. We then use a computational technique known as *Model Checking* to analyse behaviour during the crucial period between when the agent slows down in anticipation of the reward and speeds up when the reward appears.

### 2.6 Probabilistic Analysis

#### 2.6.1 Traditional behaviour based runs

We need to define a performance parameter which reflects how successful the agent has been in obtaining rewards. Since this paradigm is about waiting for delayed rewards we compare the number of successful rewards against all encounters with the landmark:

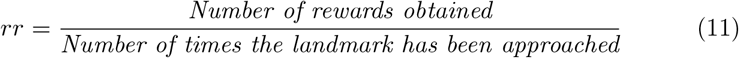

Note that this average reward is not just an academic measure but is monitored within the limbic system and then drives the levels of both serotonin and dopamine amongst others (Niv, 2007). The complete code including scripts running all scenarios are part of our open access repository (Porr et al., 2019).

Traditionally performance measures are obtained by running the experiment many times and performing a statistical analysis. They have the advantage of being close to the biological model (the behaviour of the animal) but are very time consuming to run. An alternative is model checking.

#### 2.6.2 Model checking

An alternative approach used extensively in computing science is model checking. We represent the neuronal activity and behaviour via a formal language. We then use an automatic software tool called a *model checker* to analyse our system using both simulation and verification. The model checker does this by first creating an underlying mathematical representation, which is then explored to evaluate properties. Note that, for convenience, we refer to both the formal description and the underlying mathematical representation as the *model* in this paper.

Creating a model necessarily requires us to abstract behaviour – to only contain aspects that are relevant to the properties being verified. We use model checking to focus on the core aspect of this paper, namely waiting for a reward.

The property that we want to evaluate using our model is:

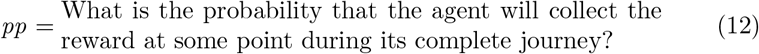

which is directly comparable to the relative reward (Eq. 11) obtained by multiple runs.

We use the model checker PRISM (Kwiatkowska et al., 2011) to determine the probabilities for our 8 scenarios from Section 2.5. PRISM is a probabilistic model checker that allows for the analysis of a number of probabilistic models including Discrete Time Markov Chains (DTMCs), Markov Decision Processes (MDPs) and Continuous Time Markov Chains (CTMCs). All of the models used in this paper are DTMCs. Models in PRISM are expressed using a high level modelling language based on the Reactive Modules formalism (Alur and Henzinger, 1999) and properties used in the verification of DTMCs are based on Probabilistic Computation Tree Logic (PCTL) (Hansson and Jonsson, 1994). In a Prism model, each module has a set of finite-valued variables which contribute to the module’s state, the global state space of the system is given by the product of the states of each module. Transitions of the model are established by way of commands, where a command consists of an (optional) action label, guard and probabilistic choice between updates. The update specifies how the variables of the module are updated when the command is executed. The probabilities for each update sum to 1. Modules interact through guards and synchronise via action labels.

As outlined above, in contrast to the behaviour based approach we focus on the crucial moment when, after learning, the agent sees the landmark, approaches it and waits in front of it to obtain the delayed reward. Our model assumes a linear search area as depicted in Fig. 6 which is our one dimensional place field from Fig. 1. Central to this is Eq. 3 which controls the speed of the agent while it approaches the landmark. The reward can then appear at any of the seven positions marked on the central line. The agent should then speed up to reach the reward. The variable *σ* is set in a way that the seven positions are spread evenly over the place field. See Porr et al. (2019) for the complete Prism code, and the appendix for the parameters.

**Figure 6:**
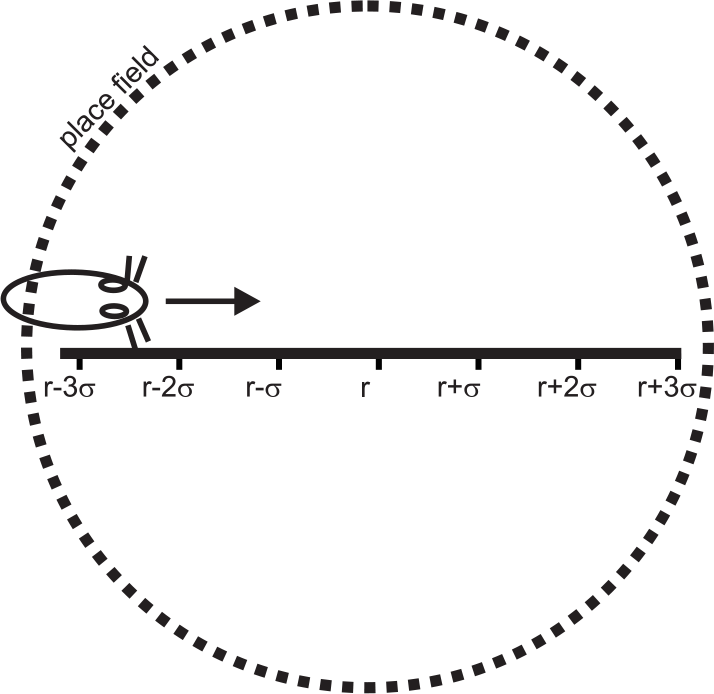
Linear representation of the behavioural experiment

To represent the behaviour of the agent and the delayed reward our model consists of two interacting modules. These are the `limbic_system` module, and the `reward_spawner` module. These modules synchronise after the agent has reached the waiting area and the `reward_spawner` delays releasing the reward. The behaviour of these modules is illustrated in Fig. 7. Note that the states in Fig. 7 actually correspond to groups of states in the underlying DTMC. The states labelled *S*_*i*_ or *t*_*j*_ in the `limbic_system` and `reward_spawner` modules correspond to all states for which variables *s* and *t* have the value *i* or *j* respectively. Note that from index 2 these states match those introduced in Fig. 1. States *S*_0_ and *S*_1_ correspond to the start of the behavioural model and a point at which random movements prior to seeing the landmark have occurred. We use the transition between the two to set `speed_type` - a variable which will contribute to the likelihood of missing the reward location later in the model.

**Figure 7:**
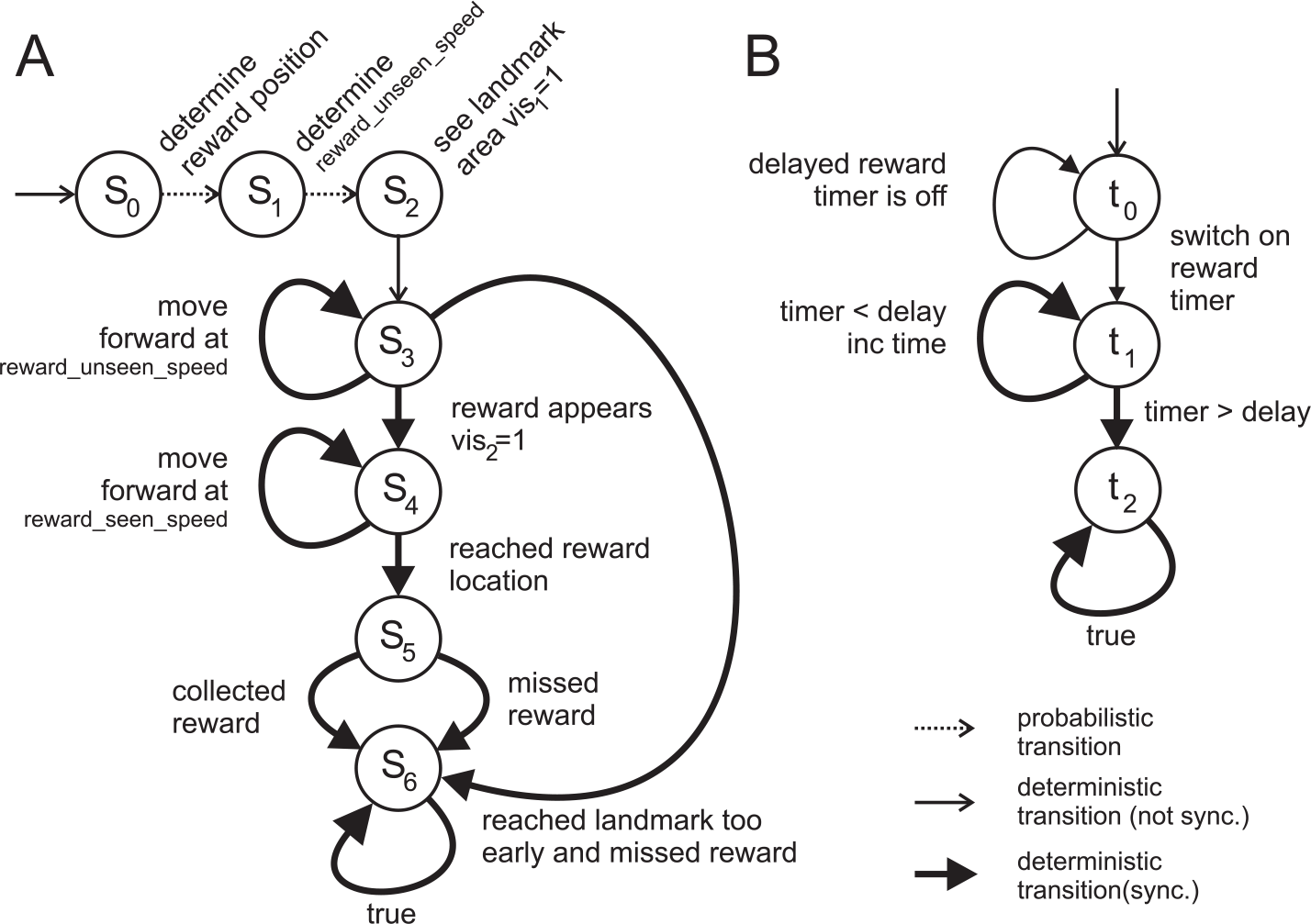
States and transitions for the Prism model

To describe our model we refer to Fig. 7. The states associated with the `limbic_system` module are described below:

*S*_0_: Initially a probabilistic choice is made as to the position of the reward (represented by the first transition in the `limbic_system` module - probabilistic choice is denoted by a dashed line). There is an equal probability of the reward appearing at each of the positions *r* + *kσ*, for *k* ∈ {−3, −2, −1, 0, 1, 2, 3} (as illustrated in Fig. 6).
*S*_1_: One of three speed types is probabilistically chosen (which will control the speed of the agent). In the behaviour-based simulation this speed variation is due to different angles at which the agent approaches the landmark and fluctuations of the weights controlling the speed. Both of these factors are abstracted to the three different speed types.
*S*_2_: The agent can see the landmark (*vis*_*1*_ = 1) and approaches it. The speed of the agent is set via Eq. 3 to: *reward_unseen_speed* = *mPFC*.
*S*_3_: The agent has reached the edge of the place field. At this point under normal conditions the serotonin concentration has increased and the agent should slow down. Different pathological conditions and/or interventions might change this and these will be the crucial part of our investigation (see Section 2.5).

At the same time as the agent enters the place field a *timer* is started which allows us to delay the release of the reward. The `reward_spawner` module then waits a predefined number of time steps (`delay`) before the reward is released. The two modules synchronise during this period (denoted by transitions with thick lines in Fig. 7), preventing the agent from speeding up until the reward has appeared.

However, if the agent is impatient and does not slow down sufficiently it will reach the (empty) landmark prematurely and miss the reward. This is reflected in the model by an immediate transition to final state *S*_6_.

*S*_4_: At this point the timer has reached the set delay time and the `reward_spawner` makes the reward appear (*vis*_*2*_ = 1) so that now both *vis*_*1*_ = 1 and *vis*_*2*_ = 1. According to Eq. 3, the speed now (*reward_seen_speed*) is set to a higher value to obtain the reward. Again, this might be compromised or improved because of pathological cases or interventions.
*S*_5_: The reward is collected if it has appeared and missed otherwise.
*S*_6_: The final state - entered whether the reward has been obtained or not.

## 3 Results

We present our results in three subsections. First we describe instructive single simulation runs which show the activities in the different nuclei and relate these to the activities. We then give statistical results from traditional behaviour based simulations followed by our model checking results.

### 3.1 Single simulation runs

In this section we show how we can use our simulation model to examine the different activities in a qualitative way to gain an intuition of the processing involved in this complex cortical and subcortical network. This is done by performing eight single instructive simulation runs according to the eight scenarios (section 2.5).

#### 3.1.1 Control run

Fig. 8 shows the signal traces of a successful run where the agent learns to approach the green landmark and to wait in front of it. The numbers in the figure correspond to those we used previously in Fig. 5. Before step 1 the agent wanders randomly. The visual signal *vis*_1,*G*_ indicates that the agent sees the green landmark. It is strongest when the agent is close to it.

1. The agent accidentally waits in front of the green landmark which then delivers food before the agent wanders off so that the agent also has a non zero *vis*_2,*G*_. At the moment the agent receives the food at 1) the VTA is triggered which then causes long term potentiation in the NAcc core so that its weight grows. This will cause the agent to approach the green landmark from a distance next time. The agent is returned to its starting point. At this point the agent also associates the place field around the green landmark with the reward which will cause a rise in the DRN activity next time and subsequent strengthening of this association. This is entirely done by the OFC which keeps track of the reward value.
2. After an unsuccessful attempt the agent sees the landmark from a distance at 2) and approaches it.
3. The agent enters the place field around the green landmark and the DRN activity rises. This now creates the crucial drop in activity in the mPFC which is caused by the activation of the 5HTR1 and 5HTR2 receptors as described in Section 2.2.2. The suppression of mPFC activity is crucial here. This can clearly be seen at the point at which the DRN activity increases. This makes the agent stop as no activity is fed downstream to the NAcc core and consequently no action is triggered. The agent waits. Any smaller distracting signals would be suppressed.
4. The reward appears and with that *vis*_2,*G*_ > 0. The overall effect is a much stronger input to the mPFC in the region of *input*_*G*_ = 2, so Eq. 3 now receives a strong input from both *vis*_1,*G*_ and *vis*_2,*G*_ which are now both amplified due to the high 5HT concentration. The high 5HT concentration makes the agent focus on the strong signal and approach the target.
5. The agent receives the food and obtains a reward. This further strengthens the association between the green landmark and the place field.

**Figure 8:**
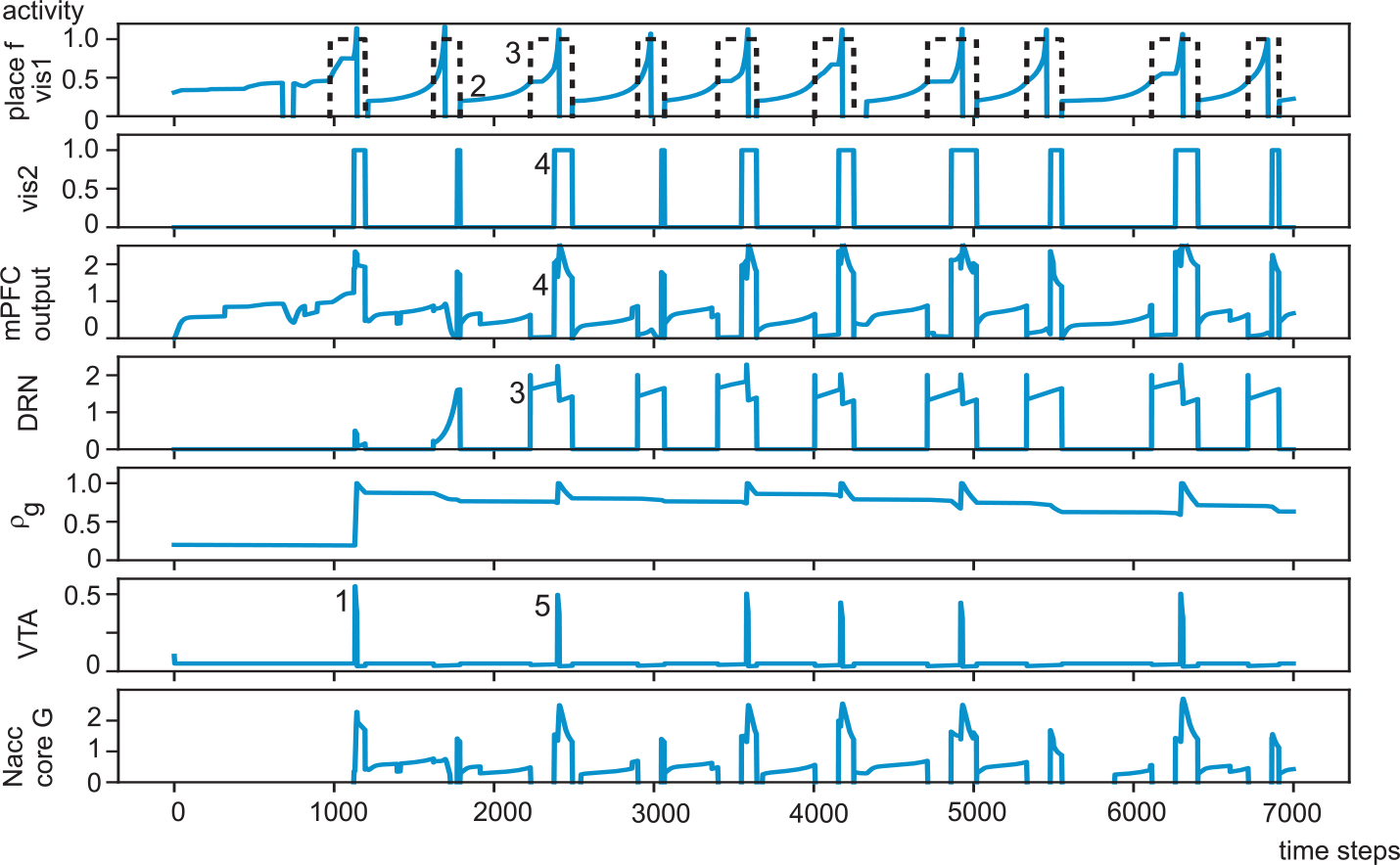
Control: Successful learning and waiting. Signal traces.

The agent is not perfect. It might miss the food because of its limited viewing angle or because it is not able to turn around quickly enough to approach the food. If this happens a negative reward prediction error is generated and the agent experiences long term depression.

Overall our simulation shows that the agent obtains rewards because 5HT causes it to wait. This is achieved by suppressing smaller signals feeding into the mPFC at high 5HT concentrations. This makes the agent wait and only approach the landmark once the additional stimulus from the food creates an overall strong and thus salient signal.

We now examine how this behaviour is altered when the 5HT is reduced and which interventions are effective.

#### 3.1.2 DRN activity reduced

Fig. 9 shows a typical run where the activity in the DRN is suppressed (i.e. *DRN_SUPP_* > 0 in Eq. 10) in addition to the inhibition from the RMTg. A division operation has been chosen because this simulates a stronger GABAergic inhibition of the DRN. Note that the most likely cause of less DRN activity is stronger GABAergic inhibition which would act as a shunting inhibition.

**Figure 9:**
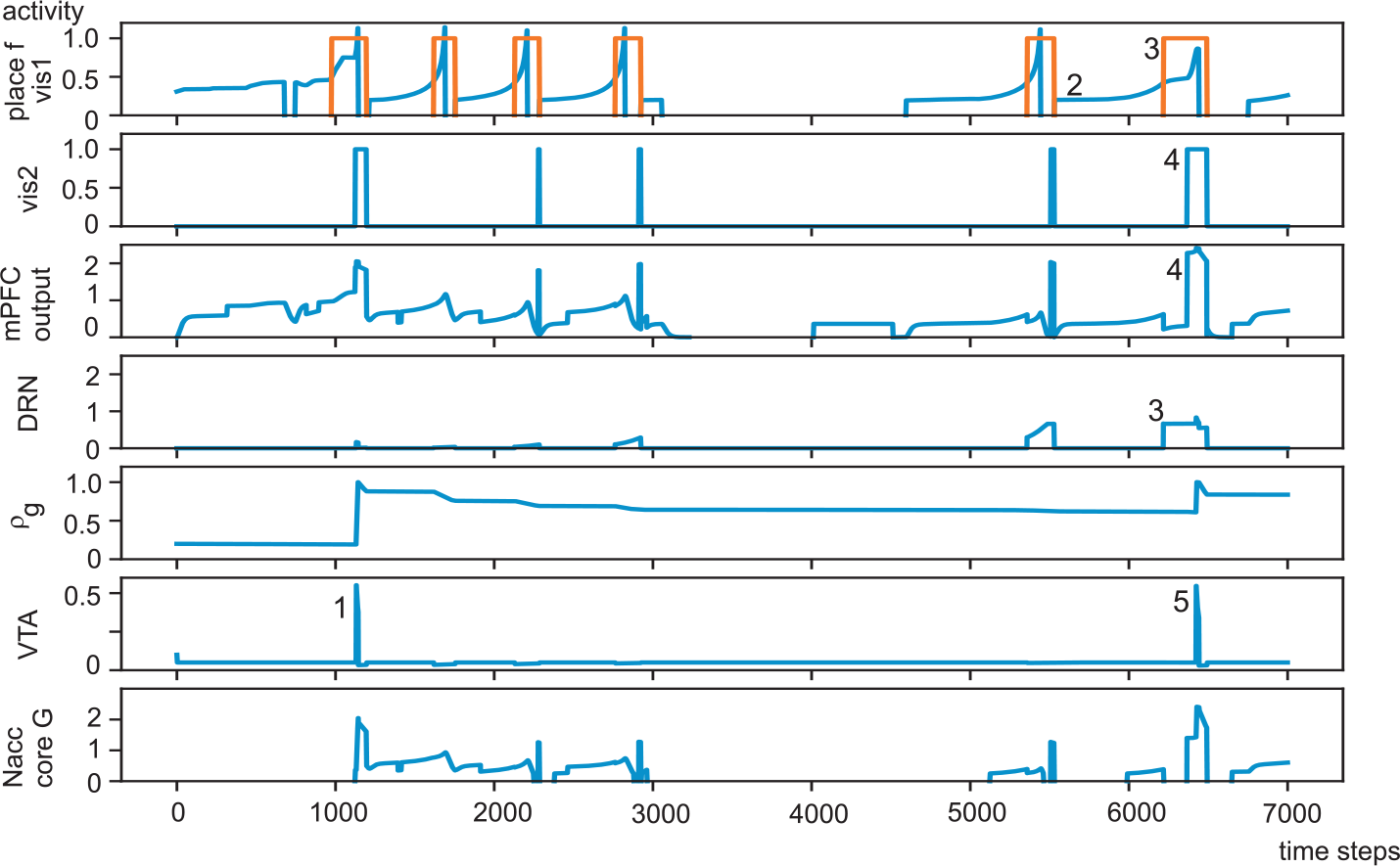
DRN activity reduced. Signal traces.

As before the the agent receives a reward at 1) but the next time it approaches the landmark it does not wait and thus does not receive a reward. This causes a negative reward prediction error and with that a decay of the weights in the NAcc. This means that the successful association with the landmark is unlearned and we see this in the decay of the weight. Thus, no waiting leads to fewer rewards and negative prediction errors which will lead to even fewer rewards in the future.

So far we have focused on sub-cortical processing. However it is well known that OFC tracks reward value as well as the NAcc shell. Indeed the OFC is possibly the more important area. We stressed earlier that this brain area is much more influenced by 5HT than by DA. Plasticity is also likely to be driven by 5HT. With reduced 5HT release plasticity changes will become slower. As a result it will be longer before the OFC learns that the area around the green landmark is potentially rewarding. This can be seen by the slow rise of the DRN activity which eventually saturates at a lower level than in the healthy condition.

In summary there are two effects caused by a depletion of 5HT:

1. Poor signalling that a reward is imminent means that the agent does not wait for the reward. This leads to fewer rewards in total.
2. Because of a lack of 5HT in the OFC plasticity is not increased when there is the potential of a reward. The OFC thus does not effectively learn the association between rewards and reward-potential cues.

#### 3.1.3 Restoring activity with SSRIs

Serotonin reuptake inhibitors (SSRIs) are important and effective drugs against depression. Fig. 10 shows the traces of a run where we have simulated the action of the SSRIs: because 5HT is not re-absorbed it continues to stimulate the receptors at a certain baseline level. We have simulated this with a bias added to the 5HT concentration (*DRN*_*OFFSET*_ > 0 in Eq. 10). In order to make it visible in the traces we have added the bias to “DRN” which is identical from the perspective of the simulation, namely that the receptors experience a constant stimulation.

**Figure 10:**
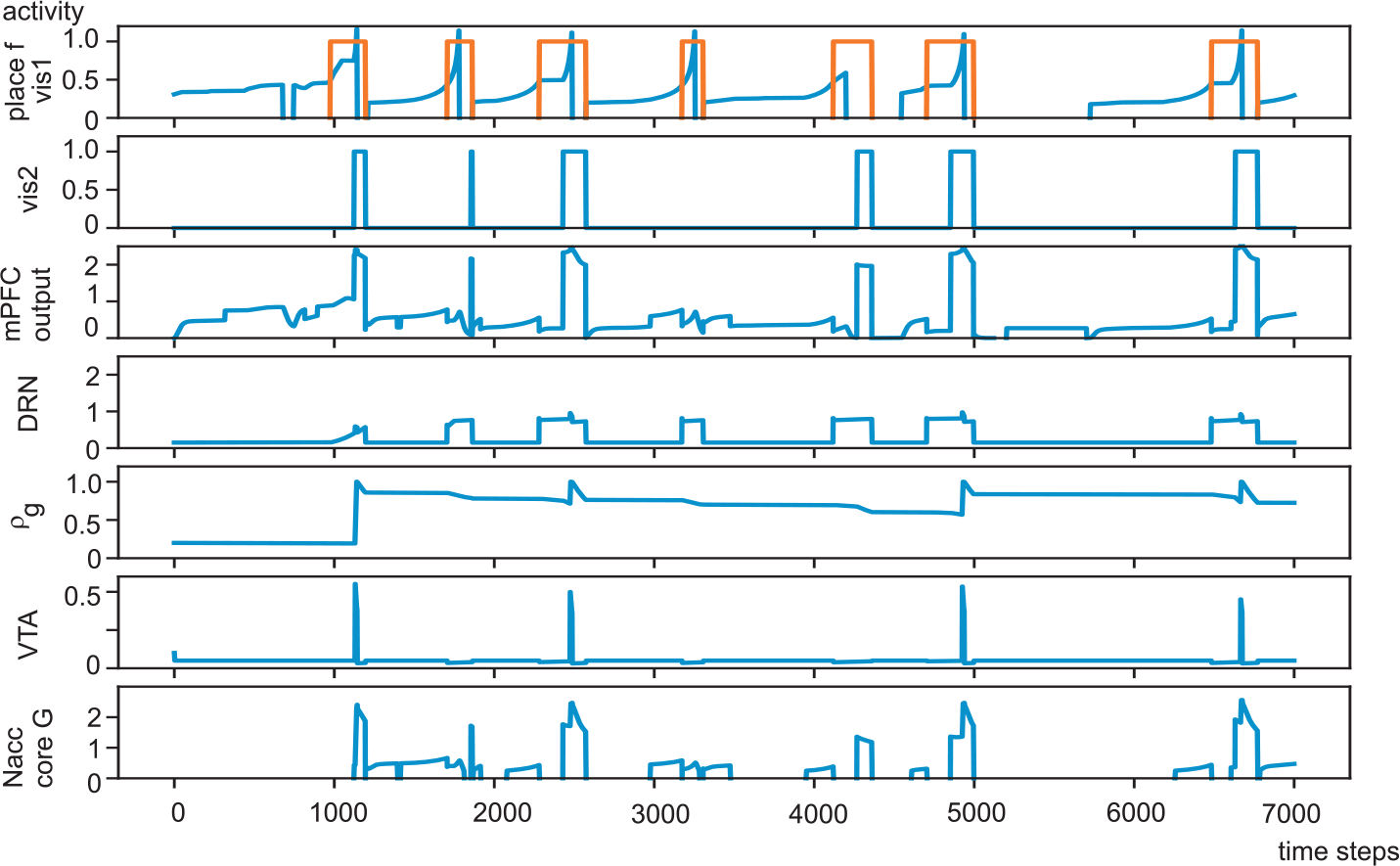
DRN activity reduced. SSRI back up. Signal traces.

The shift in the baseline has two positive effects on the learning. Learning of the reward value in the OFC is much faster because it initiates the positive feedback between OFC and DRN as soon as a reward has been triggered. We see that the increase of the DRN activity is much faster and saturates only after a few contacts with the landmark. In addition the maximum concentration of 5HT is higher which leads to the agent waiting in front of the landmark, so receiving more rewards.

In summary the SSRIs provide good relief against the problems caused by low DRN activity: enhanced plasticity in the OFC and greater reward value due to a higher 5HT signal. A point to note is that learning will become less specific due to the increase in plasticity. However, because learning is still triggered by the reward from the LH this is of minor concern.

#### 3.1.4 Restoring activity with psychedelics

Recently psychedelics have been suggested as a means to counteract the effect of loss of 5HT. Fig. 11 shows a relevant simulation run. Psychedelics particularly stimulate the 5HTR2 receptor which is responsible for the gain of the neuronal transmission. In order to understand how this is beneficial we recall the different contributions of the 5HTR1 and 5HTR2 receptors. The 5HTR1 receptor decides how small signals are to be treated. At low concentrations of 5HT small signals are amplified whereas at high 5HT concentrations they are suppressed. When the DRN is not able to release much 5HT small signals are amplified even more. This means that the agent will approach potential food sources even if their stimulus is small (and so the agent will approach any object). On the other hand the stimulation of the 5HTR2 receptor introduces a bias on the 5HTR2 receptor so that it constantly boosts the gain of the target neuron. We have simulated this by adding a constant value (*HTR2*_*OFFSET*_ > 0) to the 5HTR2 activation in Eq. 5.

**Figure 11:**
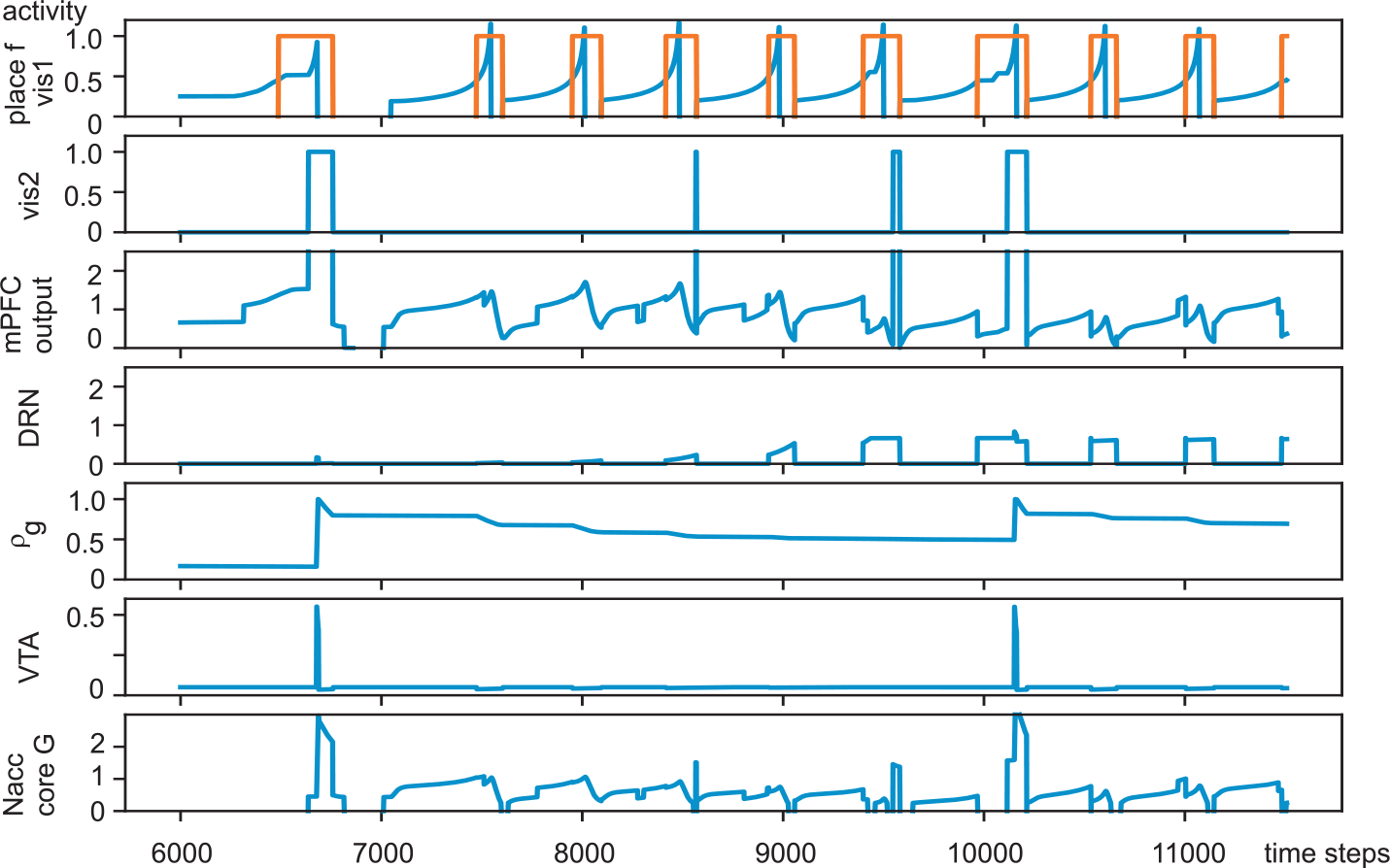
DRN activity reduced. Psychedelics. Signal traces.

Returning to Fig. 11 we can observe the effect of this bias. As the agent was attracted to the blue landmark as well as the green one it did not receive a reward until time step 6700. On the other hand, learning is probably enhanced because of higher 5HT activity. This leads to a mild beneficial effect overall and the agent eventually learns to wait in front of the landmark.

In summary the benefit of 5HTR2 agonists are in their ability to enhance exploration. This leads to more contact with the landmarks but also more disappointment.

Having developed our model and obtained intuition as to how processing in cortical and subcortical areas works, we now perform a quantitative evaluation of the overall reward obtained. First we use the simulator from the previous section by running all experiments many times. We then show the results of using model checking to investigate the crucial stage of waiting for a reward.

### 3.2 Statistical evaluation

The results of our statistical runs are shown in Fig. 12. These are ordered from bottom to top according to the 8 interventions described in Section 2.6. We have compared the results for each case using the t-test for dependent distributions which gave us significance (*p* < 0.05) for all combinations. We now describe our findings for each scenario.

**Figure 12:**
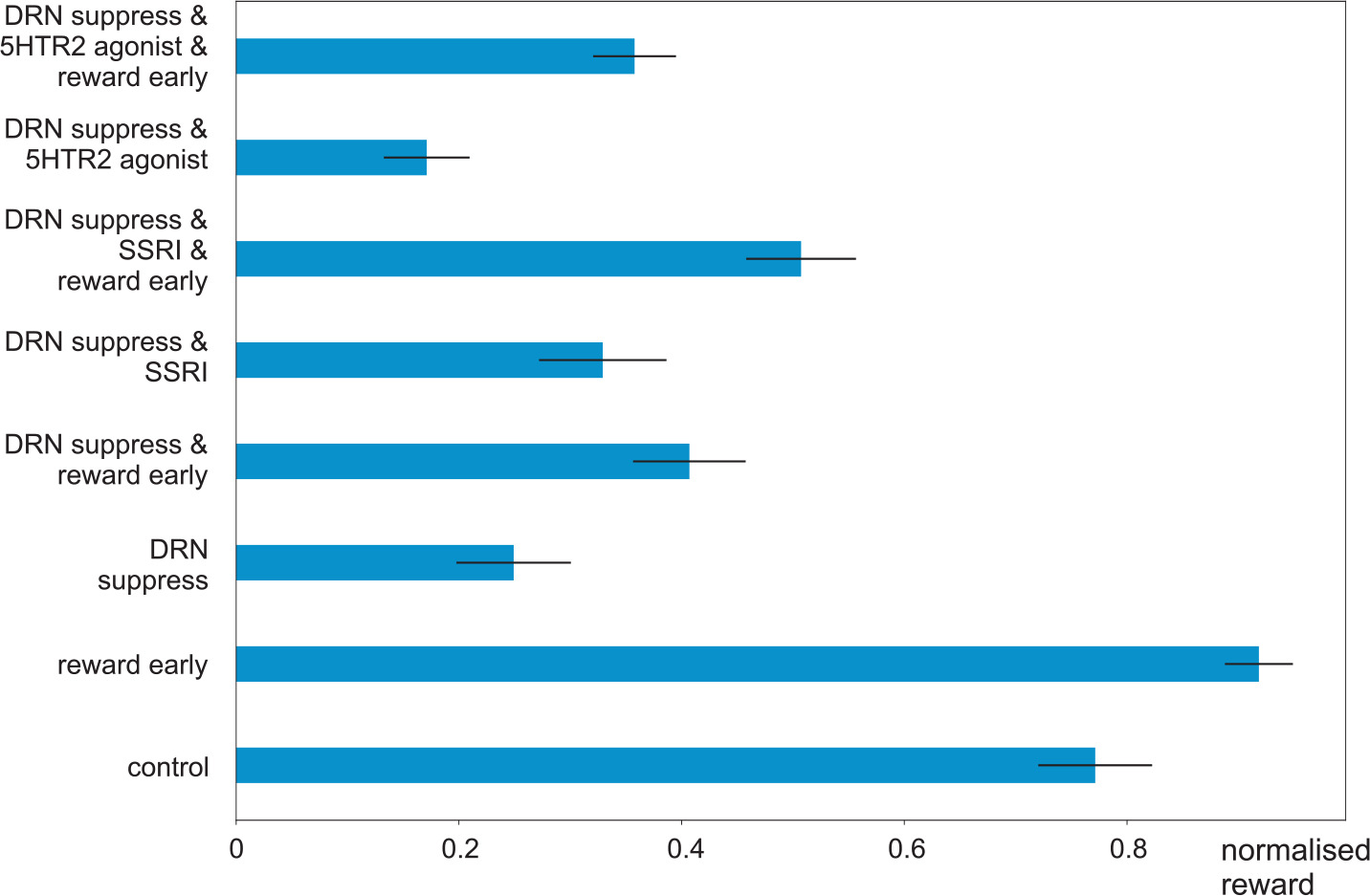
Results comparing the different scenarios

In the control condition in about 75% of all approaches to the landmark the agent receives the reward. This marginally increases when the reward is presented early.

Suppression of the DRN activity generates a very poor performance with about 25% success and a very narrow error rate. An earlier reward can improve this significantly as can the administration of SSRIs. Combining the use of SSRIs with a shorter reward delay increases performance to about 50%. This means that the agent receives the reward about half of the time. In contrast, the chance of an agent running aimlessly through the arena receiving the reward is close to zero. The improvement is still not as good as the control but has a significant improvement against the pure “DRN suppress” condition with about twice as many rewards obtained. This means that the circuits in the limbic system which track reward value will reach a higher level which in turn will feed back into the cortex.

The other approach is the use of psychedelics to increase the number of obtained rewards. In accordance with the single trial run psychedelics make it worse in this scenario where the agent needs to be patient. Stimulating the 5HTR2 receptor increases the gain of cortical processing which means that the animal becomes more impulsive and won’t wait. This leads to a significantly worse performance with 5HTR2 agonists against in particular the 3rd scenario “DRN suppress”. However, this can be significantly improved when the reward is presented earlier. This points to an interesting twist revealing under which conditions psychedelics will work: they will only work in conjunction with environmental changes - just administrating them could make the situation worse.

However the application of SSRIs plus an earlier reward is significantly better than the administration of 5HTR2 agonists in conjunction with an earlier reward.

Overall these simulations show that even a slightly earlier reward is beneficial in both cases but is essential for 5HTR2 agonists such as LSD. This is because they increase the impulsivity of the agent meaning that it cannot cope with long delayed rewards.

### 3.3 Model Checking

Fig. 13 shows the results of model checking which can be compared to those using traditional statistical methods shown in Fig. 12. Overall we observe that model checking confirms the results from the behavioural simulation where the relative rewards track closely the reward probabilities. This is particularly the case from control up to scenario 6 involving the intervention with SSRIs. However, for the scenarios involving psychedelics there is a stronger difference between the pure 5HTR2 agonist and the situation where the reward is delivered early. This means that the environmental contribution is much more emphasised during model checking. Remember that our model checking model uses a one dimensional abstraction of the behaviour so that in the case of an impatient agent there is little chance of “accidentally” waiting for the reward by detouring via intermittent distractions. Our abstraction from the behavioural model allows us to show a distinct advantage of the SSRI approach against psychedelics, at least for the delayed reward paradigm, by focusing on one key aspect – namely how 5HT changes the neuronal transfer function (Eq. 3) turning sensor stimuli into action.

**Figure 13:**
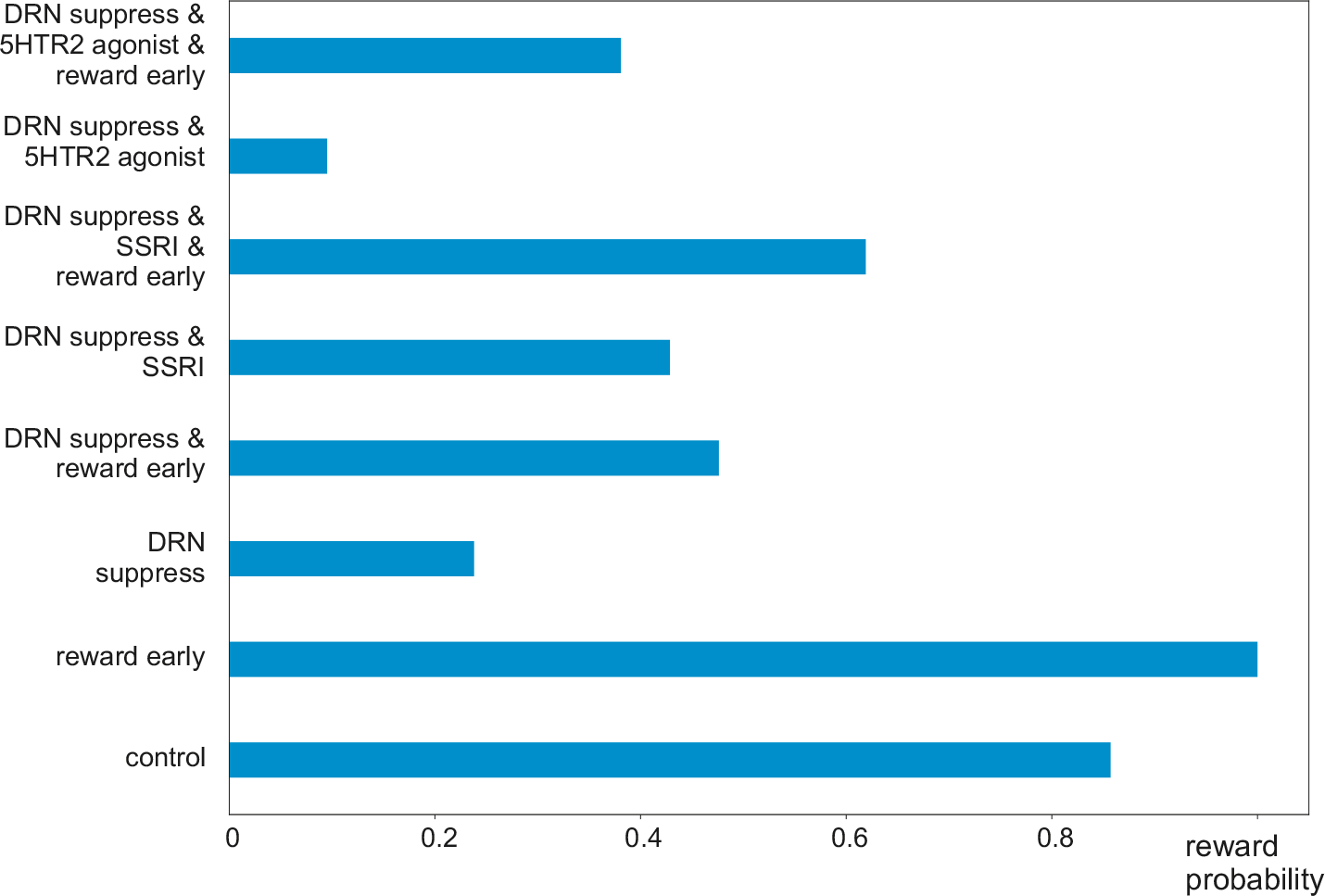
Model checking results – reward probabilities for the eight scenarios.

## 4 Discussion

We have investigated how serotonin shapes the action selection process in a simulated experiment where a rat has to wait for a delayed reward. Waiting is achieved by 5HT tuning cortical processing. At high levels of 5HT cortical processing only reacts to well learned and relevant stimuli, whereas at low levels even smaller stimuli can trigger behaviour. The pathological case of less 5HT release was then investigated with the main finding that because of less waiting the agent receives fewer rewards which in turn causes negative prediction error and eventually the unlearning of reward associations for both actions and the reward value system. This causes a downward spiral. In order to counteract this we then employed three different interventions: SSRIs, psychedelics and environmental changes making it easier to obtain the reward. Here, clearly SSRIs, environmental changes and their combination provide the best results while the introduction of psychedelics leads to mixed results.

The action of 5HT is often delayed by weeks and has been attributed to a slow de-sensitisation of, in particular, 5HTR1 and 5HTR2 receptors (Stahl, 1994). Another explanation is the ability of 5HT to boost plasticity so that neurons learn new positive associations (Scholl and Klein-Flügge, 2018). Plasticity driven by 5HT is much faster than normal and links more to ongoing behaviour. For that reason we have controlled cortical plasticity with 5HT. In contrast to intrinsic neuronal effects such as 5HT receptor de-sensitisation we argue that the slow recovery of depressed patients is because they receive more rewards causing the reward system to attach more value to sensor events. This in turn increases motivation via both the shell-vp-md-cortex pathway and an increase in tonic dopamine via the shell-vp-VTA pathway (Cofer, 1981; Dayan, 2001; Niv, 2007). We argue that improvements in the 5HT system need to filter down to the DA system and should be coupled with rewarding behaviour.

While the activity of the 5HT releasing DRN has been extensively recorded and documented (Nakamura et al., 2008; Li et al., 2016), the role of the different 5HT receptors is hotly debated. In particular the two oldest subtypes 5HT1 and 5HT2 seem to play important roles where the 5HT1 is inhibitory and the 5HT2 is excitatory (Celada et al., 2013). One might argue that these two effects cancel each other out but this is not the case: it is well established that the application of 5HT usually causes a strong depression of neuronal activity (Celada et al., 2013). This emphasises the fact that the influence of the 5HT1 receptor on signal processing is non-linear, leading to distinctly different processing according to the level of 5HT (see Eq 3).

Others argue that 5HT shifts the balance of processing because 5HT receptors are not equally distributed in the brain, and that psychedelics help to boost signal processing in the cortex because of a higher prominence of 5HTR2 receptors (Carhart-Harris et al., 2014). However, this contradicts neurophysiological findings where the cortex becomes more silent after application of 5HT (Celada et al., 2013).

Recently the role of psychedelics such as LSD as antidepressants has been widely discussed (Carhart-Harris et al., 2014; Bryson et al., 2017). The argument is that they enhance cortical processing by boosting activity in the cortex and activating 5HTR2 receptors. However, this might not always be desirable, in particular in tasks which require patience due to the fact that the activation of 5HTR2 receptors increase the gain of cortical processing (Shimegi et al., 2016). This might lead to more rewards because of more (random) activity but won’t provide measured goal directed behaviour. Psychedelics might work in situations where rewards are readily available and increased random encounters with rewards boost the reward system so increasing mood. For this reason we argue that interventions with drugs requires matching environmental interventions.

While LSD acts on the serotonergic system, NMDA receptor antagonists such as Ketamine have also shown promising results (Chaudhury et al., 2015). The positive effects of Ketamine also agree with our results in that it indirectly causes less inhibition on the Raphe Nucleus by inhibiting the activity of the Habenula so that the overall effect is disinhibitory.

In this paper we have shown how 5HT helps in the acquisition of rewards when patience is required. In our experiments 5HT made the rat focus on the relevant stimulus. A related effect would be faster response to the omission of rewards (i.e. learning to reprocess due to negative reward prediction errors (Homberg, 2012)). Reversal learning will be part of future investigations.

### A Behaviour based model

#### Lateral hypothalamus (LH)

The LH fires when a primary reward has been received.

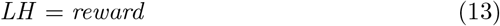

#### Ventral Tegmental Area (VTA)

The VTA receives its activity from the LH and is inhibited by the the rostromedial tegmental nucleus (RMTg).

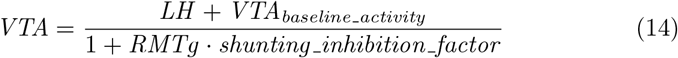

where *shunting_inhibition_factor* = 200 defines the amount of shunting inhibition on the VTA. This constant is identical for any shunting inhibition in this model. *VTA*_*baseline_activity*_ is the baseline firing rate of the VTA. At baseline neither LTP nor LTD is invoked. If the activity drops below the baseline LTD is invoked in the targets and if above baseline it is LTP.

#### Orbitofrontal Cortex (OFC)

Crucial for our model are the plastic pathways with weights 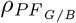 connecting the place fields (PF) to the OFC:

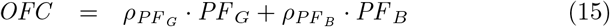

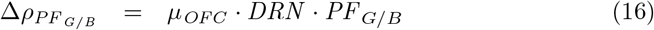

where *μ*_*OFC*_ = 0.01 is the learning rate.

#### Lateral nucleus accumbens shell

The accumbens shell also receives place field information and associates it with the help of the plastic weights 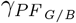:

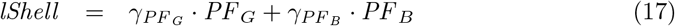

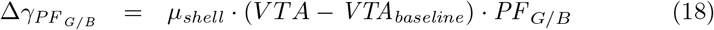

where *μ*_*shell*_ = 0.001 is the learning rate in the nucleus accumbens shell.

#### Dorsolateral ventral pallidum (dlVP)

The shell inhibits the dlVP:

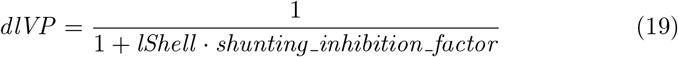

#### Entopeduncular Nucleus (EP)

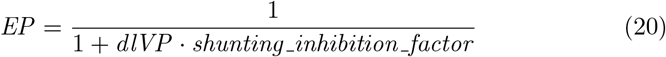

#### Lateral habenula (LHb)

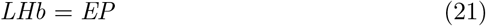

#### Rostromedial tegmental nucleus (RMTg)

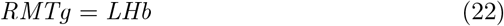

#### Nucleus Accumbens core

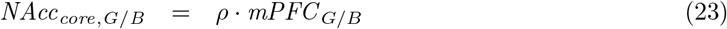

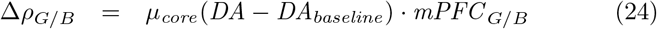

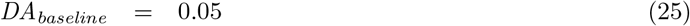

where *μ*_*core*_ = 0.075 is the learning rate in the core. The core will then disinhibit motor commands via a polysynaptic pathway involving basal ganglia structures and the motor cortex which is modeled in an abstract way. Below the agent performs exploration activity with a NAcc core activity of 0.25. Above that threshold the agent/simulated rat approaches the green or blue landmark respectively depending on which is stronger.

#### Medial Prefrontal Cortex

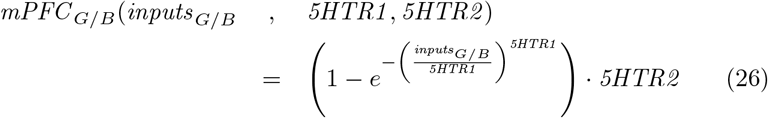

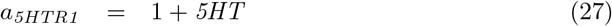

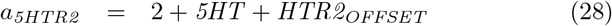

where *HTR2*_*OFFSET*_ = 0 under normal conditions and *HTR2*_*OFFSET*_ = 1 under the influence of LSD.

#### Dorsal Raphe Nucleus (DRN)

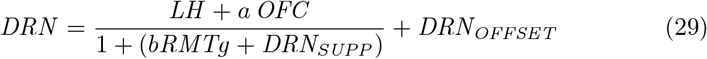

where *DRN*_*OFFSET*_ = 0 under normal conditions and *DRN*_*OFFSET*_ = 0.15 under the influence of SSRIs. The term *DRN*_*SUPP*_ = 0 under normal conditions but is *DRN*_*SUPP*_ = 4 when simulating a suppressed DRN activity due to excessive inhibition.

### B Prism model

The full Prism code is available at Porr et al. (2019).

The values of *r* and *σ* (from Fig. 6) and the speed of the agent after the reward has appeared are constants `l` and `reward_spread` which are fixed at 1000 and 330 respectively. The values of constants `delay`, `reward_unseen_speed` and `speed_uncertainty` (representing the delay in the reward appearing once the agent has reached the place field, the speed of the agent while waiting for the reward to appear, and the uncertainty in the speed - a proportion of `reward_unseen_speed`) are varied for each experiment.

The two modules `limbic_system` and `reward_spawner` synchronise after the agent has reached the waiting area and the `reward_spawner` delays releasing the reward. This is achieved in Prism via the use of *action labels*. Specifically all synchronised transitions have the action label ([timed]). This forces any such transition enabled in the `limbic_system` module to synchronise with an enabled transition in the `reward_spawner` module with the same label (if such a transition exists).

## References

Aghajanian, G. K., Sprouse, J. S., Sheldon, P., and Rasmussen, K. (1990). Electrophysiology of the central serotonin system: receptor subtypes and transducer mechanisms. Annals of the New York Academy of Sciences, 600:93–103; discussion 103.

Alur, R. and Henzinger, T. (1999). Reactive modules. FMSD, 15.

Bari, A. and Robbins, T. W. (2013). Inhibition and impulsivity: behavioral and neural basis of response control. Progress in neurobiology, 108:44–79.

Barker, E. and Blakely, R. (1995). Norepinephrine and serotonin transporters: molecular targets of antidepressant drugs. In F.E., B. and D.J., K., editors, Psychopharmacology: The Fourth Generation of Progress, pages 321–334. Raven Press.

Beckstead, R. M., Domesick, V. B., and Nauta, W. J. (1979). Efferent connections of the substantia nigra and ventral tegmental area in the rat. Brain research, 175(2):191–217.

Berthoud, H. (2004). Mind versus metabolism in the control of food intake and energy balance. Physiol Behav, 81(5):781–793.

Boureau, Y.-L. and Dayan, P. (2011). Opponency revisited: competition and cooperation between dopamine and serotonin. Neuropsychopharmacology, 36(1):74–97.

Brog, J., Salyapongse, A., Deutch, A., and Zahm, D. (1993). The patterns of afferent innervation of the core and shell in the “accumbens” part of the rat ventral striatum: immunohistochemical detection of retrogradely transported fluoro-gold. J Comp Neurol, 338(2):255–278.

Bryson, A., Carter, O., Norman, T., and Kanaan, R. (2017). 5-ht2a agonists: A novel therapy for functional neurological disorders? The international journal of neuropsychopharmacology, 20(5):422–427.

Carhart-Harris, R. L., Leech, R., Hellyer, P. J., Shanahan, M., Feilding, A., Tagliazucchi, E., Chialvo, D. R., and Nutt, D. (2014). The entropic brain: a theory of conscious states informed by neuroimaging research with psychedelic drugs. Frontiers in human neuroscience, 8:20.

Carhart-Harris, R. L. and Nutt, D. J. (2017). Serotonin and brain function: a tale of two receptors. Journal of psychopharmacology (Oxford, England), 31(9):1091–1120.

Celada, P., Puig, M. V., and Artigas, F. (2013). Serotonin modulation of cortical neurons and networks. Frontiers in integrative neuroscience, 7:25.

Chaudhury, D., Liu, H., and Han, M.-H. (2015). Neuronal correlates of depression. Cellular and molecular life sciences : CMLS, 72(24):4825–4848.

Cipriani, A., Purgato, M., Furukawa, T. A., Trespidi, C., Imperadore, G., Signoretti, A., Churchill, R., Watanabe, N., and Barbui, C. (2012). Citalopram versus other anti-depressive agents for depression. The Cochrane database of systematic reviews, (7):CD006534.

Cofer, C. N. (1981). The history of the concept of motivation. Journal of the history of the behavioral sciences, 17(1):48–53.

Cofer, C. N. and Appley, M. H. (1964). Motivation: theory and research. Wiley, New York.

Dayan, P. (2001). Motivated reinforcement learning. In Dietterich, T. G., Becker, S., and Ghahramani, Z., editors, Advances in Neural Information Processing Systems 14, Cambridge, MA. MIT Press.

Dayan, P. and Huys, Q. (2015). Serotonin’s many meanings elude simple theories. eLife, 4.

Hansson, H. and Jonsson, B. (1994). A logic for reasoning about time and reliability. Formal aspects of computing, 6(5):512–535.

Harmer, C. J. (2008). Serotonin and emotional processing: does it help explain antidepressant drug action? Neuropharmacology, 55(6):1023–1028.

Heimer, L., Zahm, D., Churchill, L., Kalivas, P., and Wohltmann, C. (1991). Specificity in the projection patterns of accumbal core and shell in the rat. Neuroscience, 41(1):89–125.

Homberg, J. R. (2012). Serotonin and decision making processes. Neuroscience and biobehavioral reviews, 36(1):218–236.

Humphries, M. D. and Prescott, T. J. (2010). The ventral basal ganglia, a selection mechanism at the crossroads of space, strategy, and reward. Progress in neurobiology, 90(4):385–417.

Kelley, A. (2004). Ventral striatal control of appetitive motivation: Role in ingestive behavior and reward-related learning. Neurosci Biobehav Rev, 27(8):765–776.

Kwiatkowska, M., Norman, G., and Parker, D. (2011). PRISM 4.0: Verification of probabilistic real-time systems. In Proc. CAV’11, LNCS 6806. Springer.

Lee, H. S., Kim, M. A., Valentino, R. J., and Waterhouse, B. D. (2003). Glutamatergic afferent projections to the dorsal raphe nucleus of the rat. Brain research, 963(1-2):57–71.

Li, Y., Zhong, W., Wang, D., Feng, Q., Liu, Z., Zhou, J., Jia, C., Hu, F., Zeng, J., Guo, Q., Fu, L., and Luo, M. (2016). Serotonin neurons in the dorsal raphe nucleus encode reward signals. Nature communications, 7:10503.

Linley, S. B., Hoover, W. B., and Vertes, R. P. (2013). Pattern of distribution of serotonergic fibers to the orbitomedial and insular cortex in the rat. Journal of chemical neuroanatomy, 48-49:29–45.

Martin-Soelch, C. (2009). Is depression associated with dysfunction of the central reward system? Biochemical Society transactions, 37(Pt 1):313–317.

Mengod, G., Cortés, R., Vilaró, M. T., and Hoyer, D. (2009). Distribution of 5-ht receptors in the central nervous system. In Jacobs, C. M. B., editor, Handbook of the Behavioral Neurobiology of Serotonin, chapter 1.6, pages 123–138. Academic Press.

Michelsen, K. A., Schmitz, C., and Steinbusch, H. W. M. (2007). The dorsal raphe nucleus–from silver stainings to a role in depression. Brain research reviews, 55(2):329–342.

Mlinar, B., Mascalchi, S., Mannaioni, G., Morini, R., and Corradetti, R. (2006). 5-ht4 receptor activation induces long-lasting epsp-spike potentiation in ca1 pyramidal neurons. The European journal of neuroscience, 24(3):719–731.

Nakamura, K., Matsumoto, M., and Hikosaka, O. (2008). Reward-dependent modulation of neuronal activity in the primate dorsal raphe nucleus. J. Neurosci., 28(20):5331–43.

Niv, Y. (2007). Cost, benefit, tonic, phasic: what do response rates tell us about dopamine and motivation? Annals of the New York Academy of Sciences, 1104:357–376.

Palacios, J. M., Waeber, C., Hoyer, D., and Mengod, G. (1990). Distribution of serotonin receptors. Annals of the New York Academy of Sciences, 600:36–52.

Peñas-Cazorla, R. and Vilarñ, M. T. (2015). Serotonin 5-ht4 receptors and forebrain cholinergic system: receptor expression in identified cell populations. Brain structure & function, 220(6):3413–3434.

Phillips, B. U., Dewan, S., Nilsson, S. R. O., Robbins, T. W., Heath, C. J., Saksida, L. M., Bussey, T. J., and Alsiö, J. (2018). Selective effects of 5-HT2C receptor modulation on performance of a novel valence-probe visual discrimination task and probabilistic reversal learning in mice. Psychopharmacology, 235(7):2101–2111.

Pollak Dorocic, I., Fürth, D., Xuan, Y., Johansson, Y., Pozzi, L., Silberberg, G., Carlén, M., and Meletis, K. (2014). A whole-brain atlas of inputs to serotonergic neurons of the dorsal and median raphe nuclei. Neuron, 83(3):663–678.

Porr, B., Trew, A., and Miller, A. (2019). An investigation into serotonergic and environmental interventions against depression in a simulated delayed reward paradigm (code). doi: https://doi.org/10.5281/zenodo.2589095.

Roberts, A. C. (2011). The importance of serotonin for orbitofrontal function. Biological psychiatry, 69(12):1185–1191.

Schildkraut, J. J. (1965). The catecholamine hypothesis of affective disorders: a review of supporting evidence. The American journal of psychiatry, 122(5):509–522.

Scholl, J. and Klein-Flügge, M. (2018). Understanding psychiatric disorder by capturing ecologically relevant features of learning and decision-making. Behavioural brain research, 355:56–75.

Schultz, W. (1998). Predictive reward signal of dopamine neurons. J Neurophysiol, 80(1):1–27.

Schultz, W., Dayan, P., and Montague, P. R. (1997). A neural substrate of prediction and reward. Science (New York, N.Y.), 275(5306):1593–1599.

Seillier, L., Lorenz, C., Kawaguchi, K., Ott, T., Nieder, A., Pourriahi, P., and Nienborg, H. (2017). Serotonin decreases the gain of visual responses in awake macaque v1. The Journal of neuroscience : the official journal of the Society for Neuroscience, 37(47):11390–11405.

Sesack, S. R. and Grace, A. A. (2010). Cortico-basal ganglia reward network: microcircuitry. Neuropsychopharmacology, 35(1):27–47.

Shimegi, S., Kimura, A., Sato, A., Aoyama, C., Mizuyama, R., Tsunoda, K., Ueda, F., Araki, S., Goya, R., and Sato, H. (2016). Cholinergic and serotonergic modulation of visual information processing in monkey v1. Journal of physiology, Paris, 110(1-2):44–51.

Stahl, S. (1994). 5ht1a receptors and pharmacotherapy. is serotonin receptor down-regulation linked to the mechanism of action of antidepressant drugs? Psychopharmacology bulletin, 30(1):39–43.

Vertes, R. P., Linley, S. B., and Hoover, W. B. (2010). Pattern of distribution of serotonergic fibers to the thalamus of the rat. Brain structure & function, 215(1):1–28.

Winter, J. C. (2009). Hallucinogens as discriminative stimuli in animals: Lsd, phenethylamines, and tryptamines. Psychopharmacology, 203(2):251–263.

Zhou, J., Jia, C., Feng, Q., Bao, J., and Luo, M. (2015). Prospective coding of dorsal raphe reward signals by the orbitofrontal cortex. The Journal of neuroscience : the official journal of the Society for Neuroscience, 35(6):2717–2730.

